# Cortical remodelling in childhood is associated with genes enriched for neurodevelopmental disorders

**DOI:** 10.1101/707042

**Authors:** G. Ball, J. Seidlitz, R. Beare, M.L. Seal

**Affiliations:** Developmental Imaging, Murdoch Children’s Research Institute, Melbourne, Australia; Developmental Neurogenomics Unit, National Institute of Mental Health, Bethesda, USA; Department of Psychiatry, University of Cambridge, Cambridge, UK; Department of Paediatrics, University of Melbourne, Melbourne, Australia

## Abstract

Cortical development during childhood and adolescence has been characterised in recent years using metrics derived from Magnetic Resonance Imaging (MRI). Changes in cortical thickness are greatest in the first two decades of life and recapitulate the genetic organisation of the cortex, highlighting the potential early impact of gene expression on differences in cortical architecture over the lifespan. It is important to further our understanding of the possible neurobiological mechanisms that underlie these changes as differences in cortical thickness may act as a potential phenotypic marker of several common neurodevelopmental and psychiatric disorders.

In this study, we combine MRI acquired from a large typically-developing childhood population (n=768) with comprehensive human gene expression databases to test the hypothesis that disrupted mechanisms common to neurodevelopmental disorders are encoded by genes expressed early in development and nested within those associated with typical cortical remodelling in childhood.

We find that differential rates of thinning across the developing cortex are associated with spatially-varying gradients of gene expression. Genes that are expressed highly in regions of accelerated thinning are expressed predominantly in cortical neurons, involved in synaptic remodeling, and associated with common cognitive and neurodevelopmental disorders. Further, we identify subsets of genes that are highly expressed in the prenatal period and jointly associated with both developmental cortical morphology and neurodevelopmental disorders.

## Introduction

The cortex is organised along a broadly hierarchical axis anchored in primary sensory regions and extending into higher-order, trans-modal cortex.^1^ This gradient is indexed by regional variations of cytoarchitecture, tissue morphology, cortical connectivity and functional network organisation^2–5^ and reflects areal patterns of cortical expansion observed across evolution and human development.^6,7^ Cortical thickness mirrors the core organisational gradient of the brain: the cortex is thinnest in primary somatosensory cortex and thickest in higher-order frontal and parietal areas.^5,8^

Cortical development during childhood and adolescence has been characterised in recent years using Magnetic Resonance Imaging (MRI). In the first two years of life, cortical thickness increases rapidly.^9^ Following this, longitudinal analyses have shown that cortical *thinning* is a prominent morphological characteristic of brain development that continues through childhood and adolescence.^10–12^ Differential rates of thinning are apparent across the cortex. In early childhood, thinning is greatest in anterior cingulate, lateral frontal and temporal cortex.^13^ Over the course of childhood and adolescence, Fjell et al.^14^ described a pattern of accelerated thinning across ventral lateral and medial frontal cortex, inferior parietal and occipital cortex, with the slowest annual change in motor and pre-motor regions and the anterior temporal lobe. Annual change in thickness is greatest in the first two decades of life and recapitulates the hierarchical, genetic organisation of the cortex, highlighting the potential early impact of gene expression on differences in cortical architecture over the lifespan.^14–16^

The biological mechanisms that underpin developmental changes to macroscopic cortical morphology are not yet clear. At a microscopic level, neuron densities can vary as much as five times across cortical areas and are inversely correlated to cortical thickness with the highest number of neurons per unit area in primary sensory regions.^4,17^ This gradient exists in contrast to the patterns of increasing neuronal soma size in frontal cortex, reflecting an increasingly complex dendritic arbor and denser intracortical connectivity with decreasing neuron density.^17,18^ In mammals, synaptic density is positively associated with cortical thickness.^19^ During human development, synaptogenesis begins during the third trimester and synaptic density increases into the second year of life before synaptic pruning results in a net elimination of synapses through to young adulthood. Although an empirical link between micro- and macro-scale cortical morphology is yet to be established, this process displays a similar heterochronicity to cortical thickness changes observed *in vivo*, with a protracted developmental trajectory in frontal cortex compared to primary auditory cortex.^20^

It is important to further our understanding of the possible neurobiological mechanisms the underlie MR-visible cortical morphology as alterations in cortical thickness could act as a potential phenotypic marker of several common neurodevelopmental and psychiatric disorders including autism spectrum disorders and schizophrenia.^21–24^ One promising approach to this problem pairs non-invasive MRI with genetic databases that detail the transcription of thousands of genes across the brain at high spatial resolution. This allows a precise comparison between patterns of gene expression and *in vivo* neuroanatomy.^25,26^ In addition, using newly available databases of single-cell transcriptomics, patterns of gene expression can be further decomposed according to cell type and developmental timing across the human lifespan.^27^ Leveraging these resources, there is emerging evidence that spatial variation of gene expression may underlie both typical variation in cortical morphometry^28,29^ and regional vulnerability to neurodevelopmental disorders.^21,30,31^

In this study, we combine MRI acquired from a typically-developing childhood population with independent post mortem gene expression databases to test the hypothesis that disrupted mechanisms common to neurodevelopmental disorders are encoded by genes expressed early in development and nested within those associated with typical cortical remodelling in childhood. We first define an imaging phenotype detailing typical patterns of cortical thinning in childhood and use this to isolate genetic pathways based on the spatial expression maps of approximately 13,000 genes in the cortex. Though comparison to independent transcriptomic datasets and prior genetic analyses, we then characterise relative gene expression according to cell-, disease- and temporal-specificity. We find that genes associated with cortical development are enriched for genes expressed in neurons, expressed highly *in utero*, and disrupted in neurodevelopmental disorders.

## Results

### Regional changes in cortical thickness over time

Using MRI data from a large, paediatric cohort^32^ we modelled regional changes in cortical thickness between ages 3 and 21 years using Gaussian Process Regression (Discovery cohort; n=537 [52.7% male], mean age [S.D.]=12.3 [5.0] years). Figure 1A shows regional change in cortical thickness over time, corrected for the effects of sex and scanner, for n=250 random cortical parcels. The majority of cortical regions (95.6%) decreased in thickness over the observation window. We calculated regional change over time relative to the rest of the cortex for each parcel (Material and Methods; Fig 1B). Cortical thinning was highest in anterior frontal and medial frontal cortex, precuneus and lateral parietal cortex and slowest in motor and premotor regions. The spatial patterning of relative cortical change over time mapped onto patterns of established functional network organisation^33^ with cortical thinning greatest in the default-mode, limbic and visual networks (Fig S1). Regional mean thickness was not correlated with relative change over time (r=−0.02, p=0.72).

**Figure 1:**
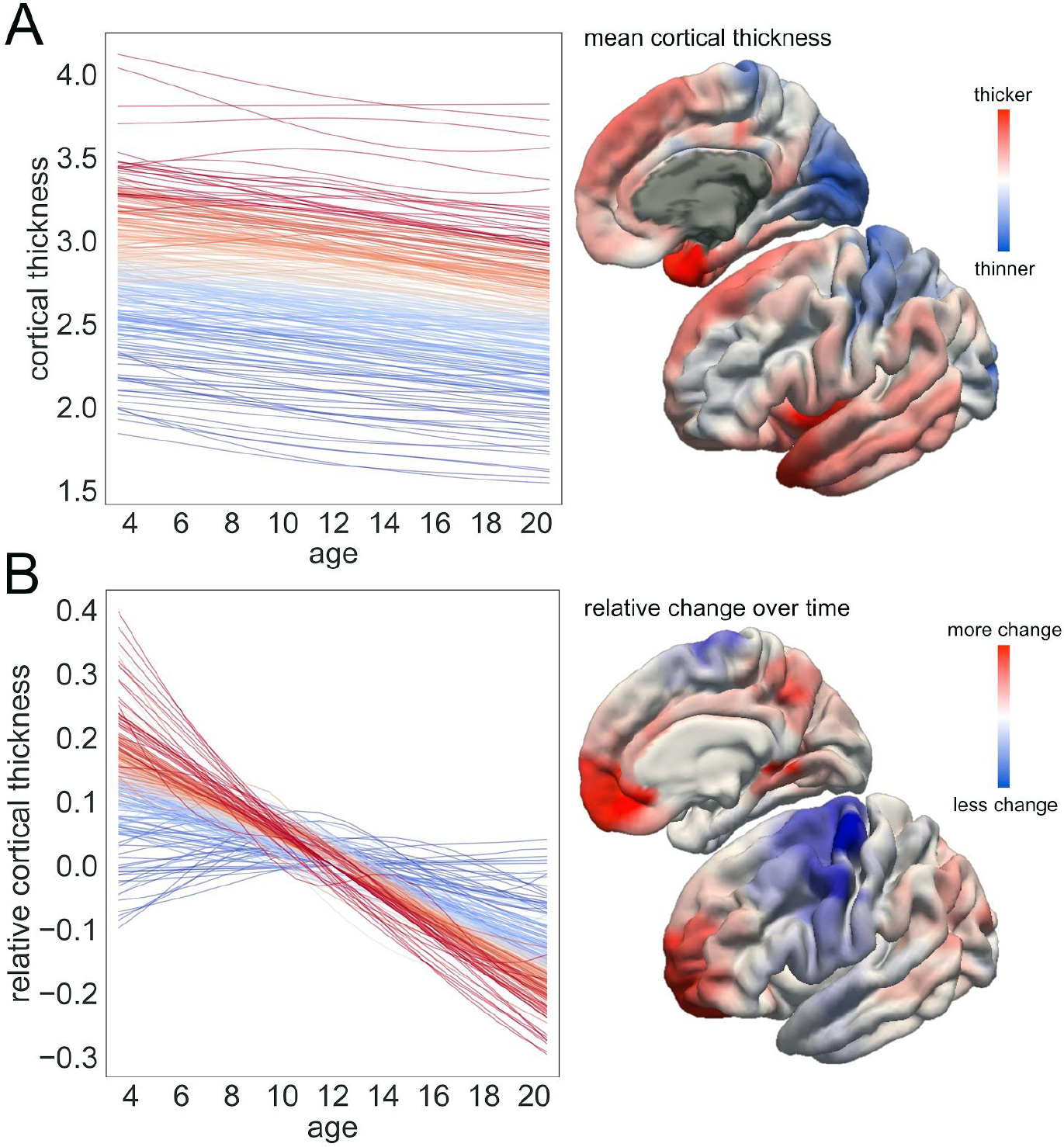
Regional changes in cortical thickness over time. Estimated trajectories are shown for each cortical parcel, coloured by mean cortical thickness (top row). Corresponding anatomical image shows cortical thickness of each parcel on the cortical surface. Bottom row shows each trajectory with the mean value removed and coloured by relative change over time. The calculation of change over time highlights regions with the greatest rate of cortical thinning. Corresponding anatomical image shows relative regional change on the cortical surface. Note: anatomical images show the lateral and medial aspects of the left hemisphere only. Surface metrics are smoothed to 5mm FWHM for visualisation.

### Cortical remodelling is associated with regional gene expression

Using relative cortical thinning as an imaging phenotype, we used Partial Least Squares Regression (PLSR) to identify a set of explanatory latent factors, or components, derived from cortical gene expression data. In our Discovery cohort, we found the first latent factor (PLS1; Figure 2) explained 41.4% of variance in our imaging phenotype. To capture the uncertainty in this estimate, we performed PLS using cortical change maps derived from 1000 randomly sampled trajectories drawn from the posterior distribution over GP functions for each region. Across all samples, the first PLS component explained between 32.2 and 42.6% (mean [S.D.]: 39.5 [0.01]%) of the variance in regional thinning. We compared this model to two null distributions, first calculating the variance explained after permuting the regional assignments of gene expression data with respect to the imaging phenotype (p<0.0001, 10,000 permutations), and second, applying random rotations to the imaging phenotype relative to gene expression data before PLS to account for the spatial autocorrelation inherent to both imaging and genetic data^34^ (p=0.005, 10,000 rotations). Using five-fold cross-validation, performing PLS using data from a subset of regions in each training fold, we confirmed that a low-rank PLS solution was optimal. Only a few factors were needed to explain our imaging phenotype. Specifying between one and ten latent factors, we found that three components together explained an average 67.2% (S.D. across folds: 3.9%) of regional variance in our imaging phenotype (46.3% [S.D. 10.4%] in unseen regions). Increasing the number of components beyond three did not significantly improve prediction of regional cortical change based on gene expression data (Fig 2A, 2B). We repeated this procedure using an alternative cortical parcellation (n=180 regions per hemisphere), and an alternative measure of change over time to demonstrate our observations were robust to experimental perturbation (Supplemental Results; Fig S2 - S3). The second and third PLS components are shown in Figure S4, from here on, we focus on the first PLS component, denoted *PLS1*, which explained the largest proportion of variance across regions corresponding to the highest correlation between regional gene expression and cortical change over time.

**Figure 2:**
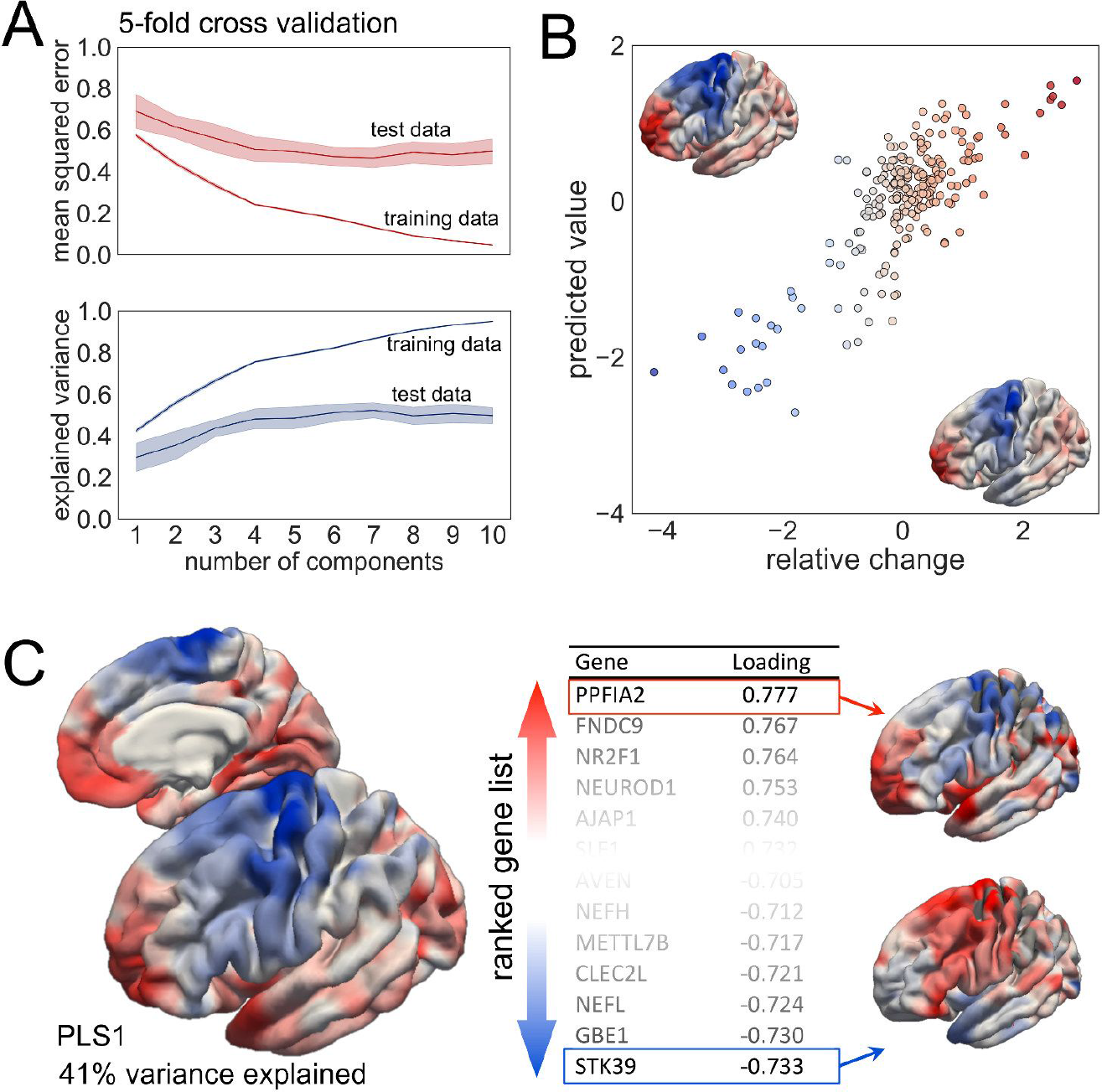
Cortical remodelling is associated with regional gene expression. A. Cross-validation results. Mean squared error (top) and variance explained (bottom) across five folds, evaluated over regions included (training data) and excluded (test data) from the PLS model. Shaded areas show 95% C.I. B. PLS results. The PLS solution with 3 components explained 64% variance in the imaging phenotype. Colour indicates relative cortical change over time. Inset shows the PLS-predicted (top corner) and original images (bottom corner). C. Component map of the first PLS component. Each component is associated with a list of genes with rank denoted by the correlation between PLS map and respective gene expression maps (inset, right). Example maps for PPFIA2 and STK39 are shown.

The PLS procedure results in a cortical map per component, representing a weighted combination of 12,799 gene expression maps derived from the AHBA data. The map associated with *PLS1* is shown in Figure 2C. Each map is associated with a list of genes, sorted by the correlation between the PLS component map and the respective maps of regional gene expression. Clear spatial correlation is evident between *PLS1* and the map of cortical change, with high expression (red areas) in the frontal and temporal poles, medial frontal cortex, and lower expression (blue) in motor and pre-motor areas. This pattern is reflected in regional expression maps of highly-ranked genes (e.g.: *PPF1A2*, *FNDC9* and *NEUROD1*) (Figure 2C). Gene loadings were highly similar across alternative cortical parcellation schemes (Spearman’s *rho*=0.84; Fig S2). The full sorted gene list is shown in Table S1.

### Highly-ranked genes are expressed in neurons and involved in synaptic organisation

Using independent gene expression data derived from five single-cell studies of the adult human cortex,^27,35–38^ we tested the hypothesis that the spatial patterning of cortical thinning was associated with regionally-variant gene expression specific to certain cell types. Cell-specific gene lists were compiled for six canonical CNS cell classes: microglia, endothelial cells, oligodendrocytes, astrocytes, excitatory and inhibitory neurons.^30^ We performed Over-Representation Analysis (ORA) for the genes ranked in the 95th centile (n=640 genes) for *PLS1* (Figure 3A). We found that genes expressed primarily in neurons, both inhibitory (fold enrichment ratio [95% C.I.]=3.36 [3.03, 3.69], hypergeometric statistic: −log10(p)=11.8, FDR corrected across cell types) and excitatory (fold enrichment=3.43 [2.97, 3.83], −log10(p)=4.75), were included in the top-ranking gene set more often than expected (Fig 3A). Genes specific to astrocytes (fold enrichment=1.40 [1.16,1.55], −log10(p)=1.29) were also over-represented within the top ranking gene list. Neuronal enrichment ratios were similar when restricting the background reference gene list (n=35,652) to only include the genes in the AHBA data that were transcribed in the cortex (n=12,799; Table S2).

**Figure 3:**
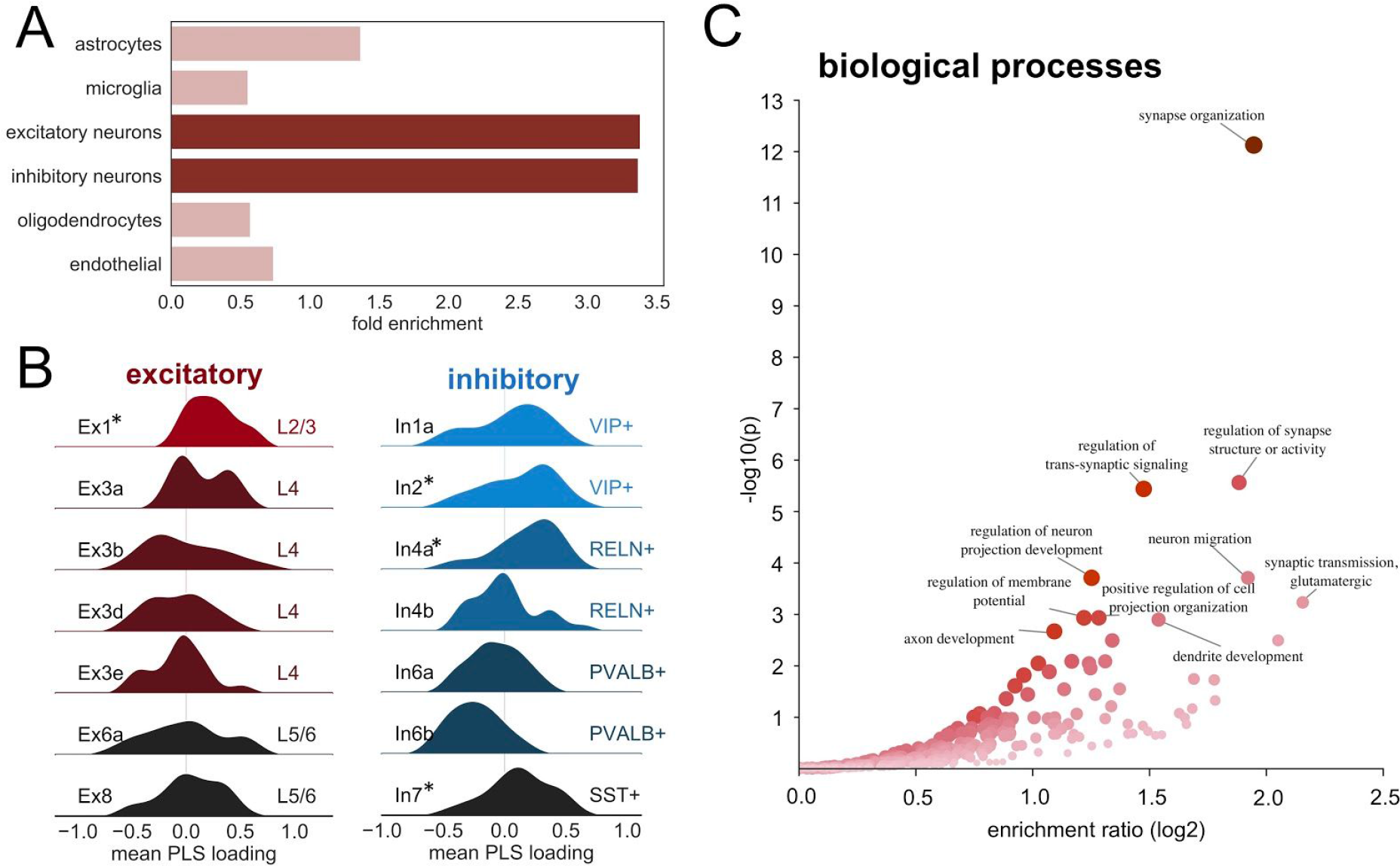
Highly-ranked genes are expressed in neurons and involved in synaptic organisation. A. Fold enrichment of six cell-specific gene lists in the top 5% of genes ranked by loading on the first PLS component. Opacity represents significant over-representation at FDR-corrected p<0.05. B. Distribution of *PLS1* gene loadings across excitatory and inhibitory neuronal subtypes.^36,39^ Colours indicate cortical layer (excitatory) or interneuron type (inhibitory). Asterisks show where gene loadings are significantly higher than expected (10,000 permutations, FDR-corrected p<0.05). C. Volcano plot showing Gene Ontology (GO) results for Biological Processes visualised with WebGestalt.^40^ Volcano plot shows enrichment of biological terms within the top ranked genes. The top 10 terms are annotated.

We performed a secondary analysis, focusing specifically on the enriched neuronal cell types, using genes expressed differentially across a set of excitatory and inhibitory neuronal subtypes.^36,39^ We identified genes expressed exclusively by each subtype, removing those with fewer than 10 exclusive genes, and tested the hypothesis that subtype-specific gene loadings on the first PLS component were significantly increased compared to randomly-selected gene sets of equal size. We found genes expressed by excitatory neurons in cortical layers 2/3 (subtype Ex1a: mean loading=0.24, FDR-corrected p=0.001) and by *VIP*+, *RELN*+ and *SST*+ inhibitory interneurons (In2: mean loading=0.13, FDR-corrected p=0.03; In4a: 0.17, p=0.03; In7: 0.11, p=0.03) localised to both upper and lower cortical layers (Figure 3B).

We reasoned that genes associated with cortical remodelling may encode important neurodevelopmental functions. To test this, we performed ORA for ontological terms associated with gene function in the top-ranked gene set. Of 640 genes, 465 (71.5%) were annotated to specific functional categories. We found a number of significantly enriched terms in the highly-ranked gene set including synaptic organisation, axonal guidance, neuron migration and axon development (FDR p<0.05) (Fig 3C; Table S3). In contrast, genes ranked in the lowest 5% were enriched for predominantly metabolic and biosynthetic processes (Figure S5).

### Genes are enriched for cognitive disorders and expressed *in utero*

To further explore putative neurodevelopmental importance of genes within the top-ranked gene set, we first tested the hypothesis that genes related to the emergence of neurodevelopmental disorders are enriched among genes associated with cortical remodelling in childhood. Using the DisGeNET database, a comprehensive catalogue of human-disease associated genes,^41^ we performed ORA of Disease Ontology (DO) terms in the top-ranked gene set. Overall, 275 of the 640 top genes (43%) were annotated to diseases or disorders in DisGeNET. Significantly enriched disease terms are shown in Figure 4A and Table S4. We found significant overlap between the top-ranked gene list and genes associated with: neurodevelopmental disorders (DO terms: Autism Spectrum Disorder/Autistic Disorder; Schizophrenia, all FDR-corrected p<0.05), mood disorders (Bipolar disorder; Major Depressive Disorder) and addiction disorders (Cocaine-Related Disorders; Chronic Alcohol Intoxication) (Fig 4B) Low ranking genes were not similarly enriched for brain-specific or cognitive disorders (Fig S5).

**Figure 4:**
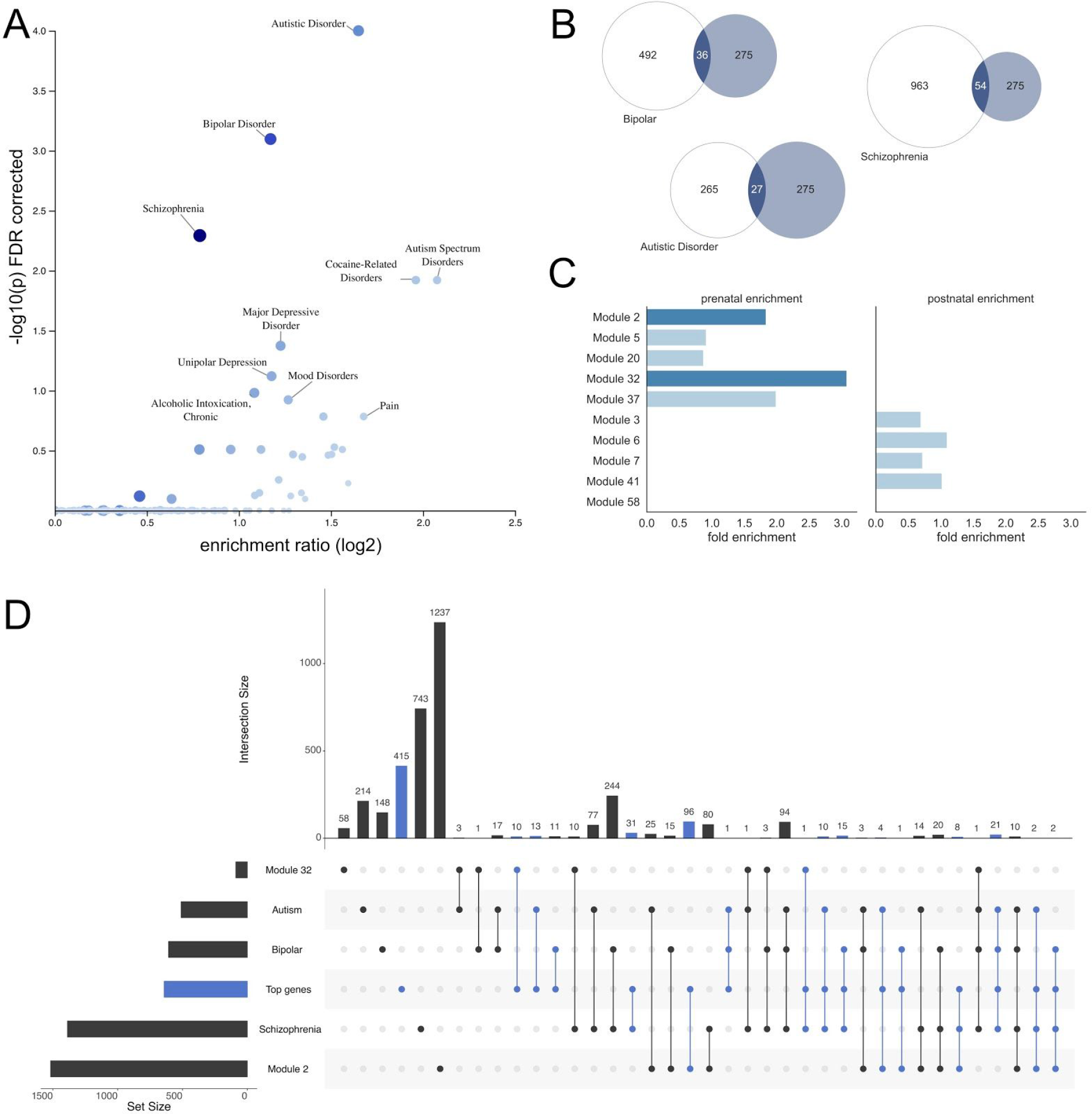
Highly-ranked genes are enriched for cognitive disorders and expressed *in utero*. A. Volcano plot shows disorders with gene sets over represented in the top ranked genes of *PLS1*. The top 10 terms are annotated. B. Venn diagrams showing the overlap between the top gene set and genes sets for Bipolar Disorder, Schizophrenia and Autistic Disorder, all p<0.05 FDR corrected. C. Gene sets associated with significant differential expression over the lifespan were defined according to Li et al.^27^. Bar chart shows fold enrichment of the genes in each module within the top-ranked gene set. Dark bars indicate FDR corrected p<0.05 (hypergeometric statistic). D. UpSet plot^42^ showing intersections of gene lists for three neurodevelopmental disorders, two prenatally expressed gene modules and the top-ranked gene list. Intersecting sets including genes in the top-ranked set are highlighted in blue.

Finally, we tested the hypothesis that genes associated with cortical remodelling in childhood and linked to neurodevelopmental disorders are expressed preferentially in early life. Using independent cortical gene expression data sampled across the lifespan, we tested for preferential enrichment of 10 gene modules with significant differential expression pre- and post-natally.^27^ Using brain-expressed genes as a reference set, we found significant enrichment of two modules (M2 and M32) in the top-ranked gene set (M2: fold enrichment = 1.53, p=6.7 × 10^−5^; M32: enrichment=2.71, p=0.02), the relative expression of both of which were higher before birth. (Fig 4C; Table S5).

To determine the overlap between genes associated with cortical remodelling, neurodevelopmental disorders and prenatal expression, we performed an intersection analysis (Fig 4D). Overall, 225 (35%) of the top-ranked genes were present in at least one other gene set. Of these, 111 were associated with at least one neurodevelopmental disorder (ASD: n=51; BPD: n=51; SCZ=81) and 124 belonged to either Modules 2 (n=113) or 32 (n=11). In total, 21 genes associated with cortical remodelling in childhood were associated with all three disorders (Table S6) and 18 genes were associated with both higher prenatal expression in Module 2 or 32 and at least one neurodevelopmental disorder (Table S7). For comparison, we also explored intersections with a non-neuronal disorder, inflammatory bowel disease (IBD; total number of genes = 564), and found 16 genes were associated with both cortical remodelling and IBD, 4 of which were also present in Module 2.

### Validation cohort

We repeated our analysis in a held-out Validation sample matched for age, sex and acqusition site (n=231, [52.4% male], mean age [S.D.]=12.3 [5.0] years). We found similar patterns of cortical thickness change over time (Spearman’s *rho*=0.68, p<0.001; Fig S6), with a strong correlation between spatial maps of the respective PLS components across samples (*rho*=0.88, p<0.001). As with the Discovery cohort, the first PLS component explained the majority of spatial variance in the imaging phenotype (PLS1: 29.6%; PLS2: 16.6%; PLS3: 10.4%) and the gene loadings on the first component were highly correlated across Discovery and Validation sets (*rho*=0.90, p<0.001). After performing over-representation analyses for gene and disease ontologies and cell types, we found similar patterns across both cohorts (Fig 5). The top-ranked genes in the validation cohort were significantly enriched for genes expressed in both inhibitory and excitatory neurons (inhibitory: fold enrichment [95% C.I.]=2.57 [2.26, 2.88], −log10(p)=2.8; excitatory: 2.67 [2.38, 2.89], 7.0).

**Figure 5:**
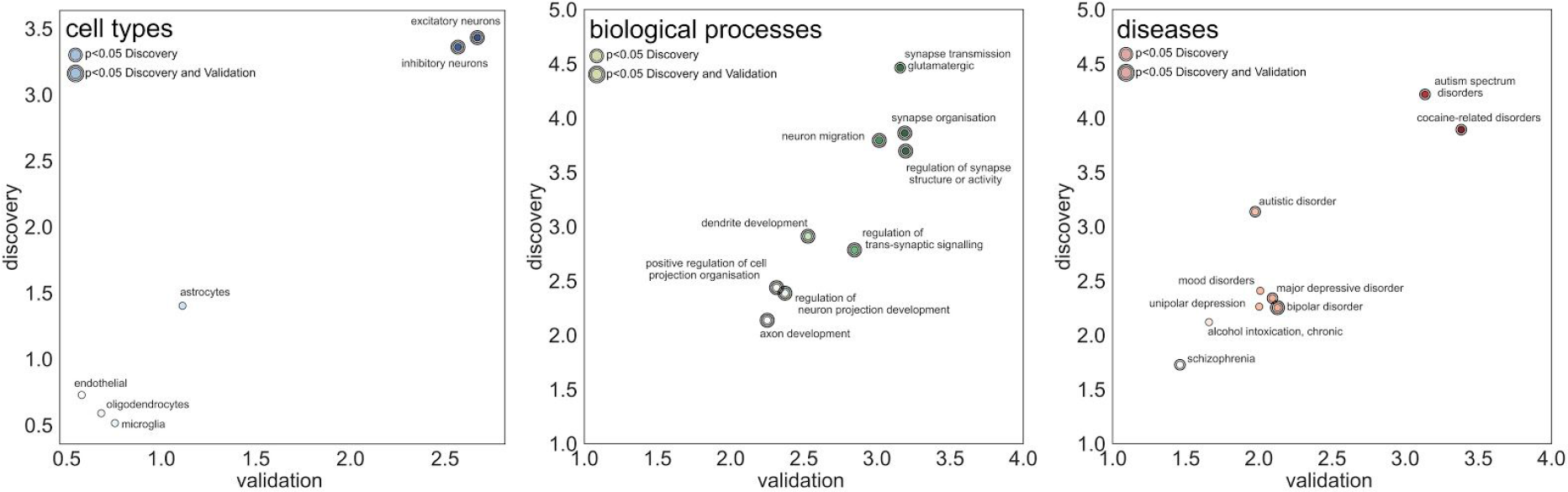
Similar patterns of gene enrichment across Discover and Validation cohorts. A. Fold enrichment ratios of cell-specific gene lists in the top 5% genes associated with *PLS1* in Discovery and Validation samples. B. Fold enrichment of biological process terms. C. Fold enrichment of disease-associated gene lists. Significance of each term at FDR-corrected p<0.05 in the Discovery cohort or in both Discovery and Validation cohort is indicated by one or two black circles, respectively.

Of the top ten significantly over-represented gene ontology terms identified in the discovery sample, nine featured in the top 20 terms in the validation sample and eight were significantly over-represented, with a high correlation between gene enrichment across samples (*rho*=0.85, p=0.003). Of the top ten disease terms in the Discovery set, nine were in the top 20 of the validation set and a similar correlation in enrichment was seen between samples (*rho*=0.70, p=0.036), although only enrichment of genes associated with bipolar disorder passed FDR correction in the validation set. Finally, we confirmed that highly-ranked genes were also enriched for gene modules that are differentially expressed over the life span with high expression *in utero* (M2: fold enrichment = 1.99, FDR-corrected p=3.2 × 10^−12^; M32: enrichment = 3.07, p=0.02).

## Discussion

We find that cortical remodelling in childhood, defined by differential rates of thinning across the developing cortex, is associated with spatially-varying gradients of gene expression. Genes that are expressed highly in regions of accelerated thinning are expressed predominantly in neurons, involved in synaptic remodeling, and associated with common cognitive and neurodevelopmental disorders. Further, we identify subsets of genes that are highly expressed in the prenatal period and jointly associated with both developmental cortical morphology and neurodevelopmental disorders. We surmise that typical patterning of cortical thinning is developmentally programmed, associated with synaptic remodelling and that phenotypic variation in neurodevelopmental and cognitive disorders may, in part, stem from disruptions to the onset of this process in the time prior to birth.

Cortical thinning is a morphological hallmark of brain development.^10,12^ In this large, typically-developing cohort spanning childhood and adolescence, we highlight a pattern of accelerated thinning in both lateral and medial aspects of the anterior frontal cortex, lateral inferior parietal cortex, and precuneus. This pattern corresponds well to previous, longitudinal studies performed across similar age ranges.^14^ As expected we found that cortical thickness decreased across the majority (96%) of the cortex between 3 and 21 years,^10^ we also found that accelerated cortical thinning is centred on regions within the default-mode network. In contrast, we find that slowest thinning is restricted to motor and pre-motor cortex, superior temporal cortex and the temporal pole. This pattern of differential thinning did not correlate with average regional cortical thickness, where the overall thinnest regions included primary somatosensory cortex and visual cortex.

In a longitudinal study of adolescents, Whitaker et al.,^28^ found that baseline cortical thickness and change in cortical thickness over time could be explained by two separable patterns of gene expression. They found that spatial variation in cortical thickness was associated with the expression of genes enriched for ontological terms encompassing metabolic and catabolic processes, and cellular responses to stimuli.^28^ This mirrors findings from an independent, cross-sectional study of adolescents demonstrating that regionally thicker cortex is associated with transcription of genes associated with astrocytes and microglia, suggesting a putative link between cortical thickness and glial associated mediation of cellular energy demands.^29,43^ We also find that gene expression in regions that remain relatively stable over time is associated with primary metabolic and energetic processes (Fig S5A). In contrast, accelerated cortical thinning over time was associated with an independent transcriptional gradient, highest in prefrontal cortex and enriched for regulation of glutamatergic signaling as well as genes specific to cortical neurons^29^ and oligodendrocytes.^28^

In this study, we confirm these observations in a larger cohort sampled over a longer time period. Our findings demonstrate that cortical remodelling captures genetic gradients associated with synaptic transmission and dendritic organisation. We extend these findings by comparison to a comprehensive database of single-cell transcriptomics studies surveying expression across six canonical cell classes. We find that accelerated cortical thinning is associated with genes expressed primarily in neurons (both inhibitory and excitatory) suggesting that cortical remodelling during childhood may, in part, depend on the dendritic remodelling and synaptic elimination processes occurring during this period.^17,18,20^

We further resolve these observations to specific cortical layers and neuronal subtypes. We found that genes expressed by excitatory neurons in layers II/III and by specific subpopulations of inhibitory neurons are highly-weighted in the composite transcriptional gradient associated with accelerated cortical thinning. Detailed characterisation of cell types in the human brain with high-throughput gene-sequencing methods has revealed several neuronal subpopulations that vary by gene expression and location in the cortex.^36,39^ By combining the spatial resolution of the AHBA with these comprehensive, single-cell databases, we are able to identify and spatially resolve specific cell populations associated with accelerated cortical thinning in development. We found that only subtype 1, located in cortical layers 2/3, showed significantly higher loading on the PLS gene component than expected by chance. Pyramidal neurons in layers 2/3 of the cortex are the last cortical layers to develop, delineating in humans at the end of the second trimester^44,45^ and form an essential component of cortical circuitry, acting as the predominant target of intra-cortical excitatory connections.^44,45^ Disruptions to the pyramidal neurons have been implicated in a number of neurodevelopmental disorders. Post mortem observations reveal a decrease in dendritic spine density localised to excitatory neurons in layer 3 of the prefrontal cortex^46^ and disruptions to layer 2/3 pyramidal neurons result in altered synaptic circuitry in mouse models of Rett syndrome, a neurological disorder with autistic-like symptoms.^47^ Inhibitory interneurons are distributed across the cortical layers, we found three subtypes associated with expression co-located with cortical thinning. Recent evidence suggests that the topographic distribution of interneuron subtypes across the cortex reflect organisational cortical gradients^48^ and both parvalbumin-positive and somatostatin-postive inhibitory interneurons have been linked to cognitive disorders including schizophrenia and bipolar disorder.^39,49,50^ The joint involvement of both excitatory and inhibitory subtypes highlights potential shared mechanisms across development and disorder centred on an (im)balance of intra-cortical excitatory/inhibitory inputs.^51,52^

In contrast to previous studies, we do not find significant enrichment of oligodendrocyte-specific gene expression (Table S2). We propose that this is likely due to differences in age between this and previous cohorts. In adolescence and adulthood, patterns of cortical thinning may best capture patterns of intracortical myelination, and thus oligodendrocyte-related gene expression, compared to at younger ages.^28,48^ This indeterminacy highlights how a number of possibly divergent, heterochronic neurobiological mechanisms may give rise to similar observed patterns of cortical morphology, as highlighted by recent reports of the dependence of MRI-based cortical thickness estimates on intracortical myelin content during development.^53,54^

By comparing genes with high expression in regions of accelerated thinning with large databases of gene-disease associations, we found significant enrichment of several cognitive disorder terms. DisGeNET is a discovery platform containing gene-disease associations compiled from multiple sources including GWAS data, animal models and literature searches.^41^ This allows a comprehensive search across >24,000 diseases and traits. Genes associated with autistic spectrum disorders were around 3 times as prevalent within genes associated with cortical remodelling than expected, a similar pattern to schizophrenia (× 1.6) and bipolar disorder (×2.25). This is supported by previous studies linking gene transcription to altered patterns of cortical thickness to autism^21^ and schizophrenia^31^ and to reports of altered cortical thickness as a common trait across multiple cognitive disorders.^24,55,56^ This suggests that neurobiological mechanisms disrupted in common cognitive diseases are shared across disorders and partly nested within those associated with typical cortical development. We derived a subset of transdiagnostic genes associated with cortical remodelling, and associated with synaptogenesis and intracortical signal transmission (Table S6). These included the neurotrophic factor *BDNF*; *SHANK3*, deletion of which results in Phelan-McDermin syndrome (22q13 deletion syndrome) characterised by intellectual disability and autistic-like traits and behaviours;^57^ as well as genes recently identified in a large-scale GWAS of schizophrenia (*GRIA1*, *CACNA1C*).^58^ This highlights how such approaches can yield biological informative insight into how disease-specific genetic alterations may lead to downstream disruptions of cortical phenotypes accessed *in vivo*.^30^

Finally, we tested the hypothesis that genes associated with cortical development are expressed most strongly prenatally. It is been noted elsewhere that a limitation of the Allen Human Brain Atlas (AHBA) data, particularly in the context of development, is the large difference between the average age of tissue donors to the gene expression database (42.5 years) and that of the experimental cohort.^28^ As such, the AHBA provides a small window on regional gene expression in adulthood with limited temporal resolution. In this study, we utilised an independent, time-resolved, brain tissue expression database to determine gene-specific expression during both pre- and post-natal life.^27^ Inter-regional differences in cortical gene expression are maximal during the late fetal period, with expression across regions becoming more similar over childhood and adolescence.^27,59^ Additionally, a number of genes display differential expression across the lifespan with a marked transition period around the time of birth. Li et al. defined a set of modules with maximal expression before or after birth.^27^ We found that prenatal expression was enriched among genes associated with cortical remodelling in childhood, establishing a putative temporal ordering between genetic and morphometric processes. This supports the hypothesis that transcriptional diversity across the cortex underlies morphometric patterning assessed with MRI, is developmentally programmed and, if disrupted, can lead to phenotypic variation observed in common neurocognitive disorders.

There are a number of limitations to this study. Although the cohort we analysed was large, enabling us to perform an internal validation of observations, our conclusions would be strengthened by longitudinal observations allowing an improved characterisation of developmental trajectories.^60^ As is common with other imaging-transcriptomic studies, we clarify that our observations are correlational in nature: the genetic and imaging data is derived from separate cohorts and as such our conclusions require additional empirical validation, in animal models for example. Finally, we find that genes associated with cortical remodelling are associated with a number of cognitive disorders. We note that, although significant, this enrichment only relates to a small subset of disease-associated genes in each case (Fig 4B). It is unlikely that a single imaging phenotype would capture the full range of diverse aetiologies that likely underpin these disorders, nor the heterogenous phenotypes observed across clinical populations. These findings represent one such possible pathway linking potential neurobiological mechanisms across typical and atypical cortical development.

In summary, we define a pattern of typical cortical development in childhood that aligns with spatially-varying genetic gradients that are associated with synaptic remodelling in neurons and potentially disrupted in common cognitive disorders.

## Materials and Methods

### Cohort

Data were acquired from the Pediatric Imaging, Neurocognition and Genetics (PING) Study.^32^ This cohort comprises a large, typically-developing population with participants from several US sites included across a wide age and socioeconomic range. The human research protections programs and institutional review boards at all institutions participating in the PING study approved all experimental and consenting procedures, and all methods were performed in accordance with the relevant guidelines and regulations.^61^ Written parental informed consent was obtained for all PING subjects below the age of 18 and directly from all participants aged 18 years or older. Exclusion criteria included: a) neurological disorders; b) history of head trauma; c) preterm birth (less than 36 weeks); d) diagnosis of an autism spectrum disorder, bipolar disorder, schizophrenia, or mental retardation; e) pregnancy; and f) daily illicit drug use by the mother for more than one trimester.^32^ Data from the PING study are made available via the NIMH Data Archive (https://nda.nih.gov/edit_collection.html?id=2607).

The PING cohort included 1493 participants aged 3 to 21 years, of whom 1249 also had neuroimaging data. Of these, n=773 were available to download (March 2016). After quality control and image processing (see below), the final cohort comprised n=768 participants (mean [S.D] age = 12.28 {5.02] years; 404 male) acquired from seven study scanners/sites. Site/scanner-specific demographic data are shown in Table S8.

The full cohort was randomly split into **Discovery** (n=537) and **Validation** cohorts (n=231), ensuring that the distributions of age, sex and site within the subsets matched the original cohort (Table S8). All analyses were performed first in the Discovery cohort, with the second cohort acting as independent validation of our observations.

### Neuroimaging data

3 Tesla, T1-weighted images were acquired using standardized high-resolution 3D RF-spoiled gradient echo sequence with prospective motion correction (PROMO), with pulse sequences optimized for equivalence in contrast properties across scanner manufacturers (GE, Siemens, and Phillips) and models (for details, see Jernigan et al.)^32^ Site-specific imaging parameters are shown in Table S9.

### Image processing

Structural T1 volumes were processed as described previously.^62^ Briefly, vertex-wise maps of cortical thickness were constructed with FreeSurfer 5.3 (http://surfer.nmr.mgh.harvard.edu). This process includes removal of non-brain tissue, transformation to Talairach space, intensity normalisation, tissue segmentation and tessellation of the grey matter/white matter boundary followed by automated topology correction. Cortical geometry was matched across individual surfaces using spherical registration and maps smoothed to 10mm FWHM^8,63–65^ To reduce computational load prior to modelling, cortical thickness data were downsampled to the *fsaverage5* surface comprising 10242 vertices per hemisphere. Due to the asymmetric tissue sampling of the Allen Human Brain Atlas (see below), all analysis was performed in the left cortical hemisphere only.

Quality control assessment for the PING data is detailed in Jernigan et al.^32^ Images were inspected for excessive distortion, operator compliance, or scanner malfunction. Specifically, T1-weighted images were examined slice-by-slice for evidence of motion artefacts or ghosting and rated as acceptable, or recommended for re-scanning. We performed additional, on-site, visual quality control.^62^ Any images that failed initial surface reconstruction, or returned surfaces with topological errors, were manually checked and re-submitted to FreeSurfer.

### Modelling regional cortical trajectories

For all subjects, cortical thickness maps were parcellated into n=250 regions per hemisphere with approximately equal area.^66^ Regional thickness was calculated as the 95% trimmed mean of all vertex values containing within the assigned region. We repeated our analysis using an alternative parcellation scheme comprising n=180 labels per hemisphere based on cortical anatomy (HCP; Fig S1).^67^

For each region, we used Gaussian Process Regression (GPR) to model the relationship between each individual’s age and cortical thickness, including sex and study site as covariates. GPR is a flexible, Bayesian modelling approach for the probabilistic prediction of continuous variables without the need to specify the relationship’s functional form (linear, quadratic, etc.). We briefly review GPR below, for a technical overview please refer to Rasmussen & Williams.^68^ This approach has been used successfully to model relationships between age and MRI-derived metrics in previous studies^69,70^ and has been shown to ably handle multi-site MRI data.^71^

A Gaussian Process is a distribution of functions that map *x* → *y* and is fully specified by a mean function, *m* = μ(*x*) = 0, and a covariance function, *cov* = *k*(*x*, *x*′), such that *f* ~ *GP*(*m*, *cov*).^68^ The covariance function, *k*, models the dependence of values of *f* at different values of *x*, *x*_*i*_ and *x*_*j*_, respectively. Here, we define *k*(*x*_*i*_, *x*_*j*_) as:

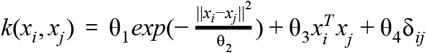

where, δ = *noise level if i* ≠ *j*, *else* 0, modelling observation noise, and θ_1,…,4_ are hyperparameters learned from the data by maximising log-marginal-likelihood. We implement GPR using *scikit-learn* (v0.20.0) in Python (v3.6.8).

For each region, we obtain predicted age trajectories, corrected for sex and imaging site, at 18 equally-spaced points between 3.5 and 20.5 years (Fig 1). To account for variance in the model, we generate 1000 randomly sampled, alternative trajectories from the posterior distribution over the function, *f*, for each region. GPR was performed separately in the *Discovery* and *Validation* cohorts.

### Cortical remodelling

To isolate regional changes in cortical thickness, we performed a non-parametric assessment of relative change over time. Using cortical trajectories estimated using GPR, we ranked each region based on relative thickness at either end of the observed timeframe (3.5 and 20.5 years), such that regions that are relatively thinner compared to the rest of the cortex had a higher rank. This rank transformation accounts for both non-normally distributed regional thicknesses at each timepoint and differences in mean thickness over time, ensuring that the non-parametric slope estimate is not dependent on initial, or final, cortical thickness. We then estimated the ‘rank change’ of each region between 3.5 and 20.5 years: *rank change* = *rank*_2_ − *rank*_1_. As such, positive values reflect areas that are become relatively thinner over time in comparison to other cortical regions (i.e.: the regional rank becomes higher over time). Finally, relative change in rank is Z-scored prior to further analysis. We repeated our analysis using an alternative definition of change over time: calculating the raw slope between estimates of cortical thickness at the beginning and end of our observation window before Z-score normalisation, although this did not affect our reported observations (Supplemental Results; Fig S2).

### Transcriptomic data

Bulk tissue microarray data were acquired from the Allen Human Brain Atlas (AHBA), a high resolution transcriptomic database comprising expression measures for more than 20,000 genes taken from 3702 spatially distinct tissue samples from six post-mortem brains.^14^

We employed a recently published pipeline to estimate regional gene expression levels across the cortex.^66^ Initially, genes were assigned to the AHBA probe set using Re-Annotator (January 2019),^72^ resulting in a probe set corresponding to 20,250 unique genes. For each brain, samples from the brainstem and cerebellum were removed and probes excluded if a) expression levels did not exceed background noise in more than 10% of cortical samples and b) demonstrated a low correlation (Spearman *rho* < 0.1) to reference RNA-seq data acquired from the same tissue.^73^ Samples were assigned to cortical regions using the same parcellation scheme applied to the cortical thickness data, with a maximum distance threshold of 2mm.^66,74^ Expression data were normalised within subject with an outlier-robust sigmoid function and averaged across subjects within each cortical region. Of the six AHBA samples, only two have microarray data from the right hemisphere. Due to this sparse sampling, we used only data from the left cortical hemisphere for analysis. Cortical regions that did not contain any samples were also removed (n=23). This process resulted in measurement of regional expression for 12,799 genes across 237 regions of the left cortex. As with the imaging data, regional gene expression data were also derived for the alternative HCP cortical parcellation (n=177/180 regions with samples in left hemisphere).

Code to perform this pipeline is available at: https://github.com/BMHLab/AHBAprocessing. The processing options used in this study are listed in Table S10.

### Partial least squares regression

Partial least squares regression (PLSR) is a multivariate extension of ordinary least squares linear regression that is well-suited to high-dimensional datasets such as neuroimaging and genetic data. PLSR works under the assumption that the dependent variable can be explained by a set of latent factors or components, each a linear combination of the predictor, or independent, variables. We used the SIMPLS algorithm^75^ to estimate weight vectors, *w* and *q*, such that:

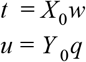

Where *X*_0_ and *Y* _0_ represent the centred *n*-region by *m*-gene regional expression data matrix and the centred *n*-region imaging phenotype, respectively, and *t* and *u* represent the *X* and *Y* component *scores*. This iterative procedure is followed by a matrix deflation step, before the estimation of a second, third, and so on, orthogonal sets of component scores (*t*_1_…*t*_*n*_, *u*_1_…*u*_*n*_). For each component, the corresponding weights are calculated to ensure maximum covariance between *t* and *u* and the component *loadings* are calculated by regressing *X*_0_ and *Y* _0_ against *t*_*n*_. After calculating scores for each PLS component, we normalise the columns of *X* and *Y* to unit norm such that the factor loadings, *X* · *t* and *Y* · *t*, are equivalent to correlations and are bounded by −1 and 1.

Thus, the PLSR results in a set of components, each associated with a *score*, defined as the weighted combination of gene expression profiles for each region, and a set of *loadings*, defined as the correlation between each gene’s expression profile and the respective component score. For each component, we sort genes according to their loading, such that genes with a high positive loading have relatively higher gene expression in cortical regions that change most over childhood and lower expression in regions that remain stable over time.

We performed five-fold cross-validation to estimate the optimal number of factors required to explain variance in PLS. For each fold, we excluded data from 20% of cortical regions before performing PLS, then estimated the mean squared error and % explained variance for the left-out regions using the estimated PLS model, before repeating with the next fold. We performed PLS with an increasing number of specified components from 1 to 20.

Statistical significance of the PLS model was determined using two non-parametric methods. For the first, we randomly permuted the regional assignment of gene expression data with respect to the imaging phenotype 10,000 times, each time calculating the variance explained in the imaging data. For the second, to account for the spatial autocorrelation inherent to both the imaging and genetic data, we applied 10,000 random rotations to the regional cortical change data before recalculating cortical change within the original cortical parcels and performing PLS.^76,77^ Unlike standard permutation procedures, this spatially-constrained permutation accounts for similarity, and therefore non-independence, between neighbouring regions.^34^

### Gene enrichment

We performed Over-Representation Analysis (ORA) for Gene Ontology (GO) and Disease Ontology (DO) terms using WebGestalt.^40^ Genes with component loadings in the 95th centile for the first PLS component were selected and compared to a whole genome reference set defined by the Agilent 8×60k microarray. The final list of 12,799 brain-expressed genes were also used as an alternative reference set (Tables S2-S5). We also used an alternative approach, Gene Set Enrichment Analysis (GSEA), that does not require an *a priori* threshold to be set to select genes of interest (Tables S11 and S12).

### Cell-specific gene expression

To determine the cell-specific expression of selected genes, we compiled data from five different single-cell studies using postmortem cortical samples in human postnatal subjects.^27,35–38^ We clustered the study-specific gene sets into canonical cell classes as described previously.^30^ This resulted in a set of six cell-specific gene lists, one for each cell class: astroctyes, endothelial cells, excitatory neurons, inhibitory neurons and oligodendrocytes.

We performed ORA for each cell-specific gene list, calculating the hypergeometric statistic as:

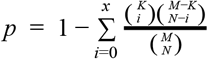

Where *p* is the probability of finding *x* or more genes from a cell-specific gene list *K* in a set of randomly selected genes, *N*, drawn from a reference set, *M*. Fold enrichment ratios were calculated as the proportion of cell-specific genes in the top-ranked list, compared to the proportion in the full reference set and statistics were FDR-corrected across cell classes.

Additional gene lists were obtained from excitatory and inhibitory neuronal subtypes,^36^ 10 modules defined in Li et al with significant patterns of differential expression over the lifespan^27^ and from disease-association lists for DO terms: Autistic disorder, Bipolar disorder and Schizophrenia in DisGeNET.^41^

## Acknowledgments

This research was conducted within the Developmental Imaging research group, Murdoch Childrens Research Institute and the Children’s MRI Centre, Royal Children’s Hospital, Melbourne, Victoria. It was supported by the Murdoch Childrens Research Institute, the Royal Children’s Hospital, Department of Paediatrics, The University of Melbourne and the Victorian Government’s Operational Infrastructure Support Program. The project was generously supported by RCH1000, a unique arm of The Royal Children’s Hospital Foundation devoted to raising funds for research at The Royal Children’s Hospital.

Data and/or research tools used in the preparation of this manuscript were obtained and analyzed from the controlled access datasets distributed from the NIMH-supported Research Domain Criteria Database (RDoCdb). RDoCdb is a collaborative informatics system created by the National Institute of Mental Health to store and share data resulting from grants funded through the Research Domain Criteria (RDoC) project. Dataset identifier(s): [DOI: 10.15154/1504093].

